# *Laccaria bicolor* MiSSP8 is a small-secreted protein decisive for the establishment of the ectomycorrhizal symbiosis

**DOI:** 10.1101/218131

**Authors:** Clément Pellegrin, Yohann Daguerre, Joske Ruytinx, Frédéric Guinet, Minna Kemppainen, Nicolas Frei dit Frey, Virginie Puech-Pagès, Arnaud Hecker, Alejandro G. Pardo, Francis M. Martin, Claire Veneault-Fourrey

## Abstract

The ectomycorrhizal symbiosis is a predominant tree-microbe interaction in forest ecosystems sustaining tree growth and health. Its establishment and functioning implies a long-term and intimate relationship between the soil-borne fungi and the roots of trees. Mycorrhiza-induced Small Secreted Proteins (MiSSPs) are hypothesized as keystone symbiotic proteins, required to set up the symbiosis by modifying the host metabolism and/or building the symbiotic interfaces.

*L. bicolor MiSSP8* is the third most highly induced MiSSPs in symbiotic tissues and it is also expressed in fruiting bodies. The *MiSSP8-*RNAi knockdown mutants are strongly impaired in their mycorrhization ability with *Populus*, with the lack of fungal mantle and Hartig net development due to a lack of hyphal aggregation. MiSSP8 C-terminus displays a repetitive motif containing a kexin cleavage site, recognized by KEX2 *in vitro*. This suggests MiSSP8 protein might be cleaved into small peptides. Moreover, the MiSSP8 repetitive motif is found in other proteins predicted secreted by both saprotrophic and ectomycorrhizal fungi. Thus, our data indicate that MiSSP8 is a small-secreted protein involved at early stages of ectomycorrhizal symbiosis, likely by regulating hyphal aggregation and pseudoparenchyma formation.

## Introduction

Forest soils contain a wide diversity of microorganisms displaying multiple nutrition modes from saprotrophy to pathogenicity, through mutualism (Buée *et al.*, 2009; Uroz *et al.*, 2010; Fierer *et al.*, 2007). Tree-associated microbes are considered as key drivers of tree health, productivity, and ecosystem functionality (Berg *et al.*, 2014; 2015). In particular, fungal communities are primary contributors to carbon and nitrogen cycling in forest ecosystems (van der Hejden, 1998; Lindahl and Tunlid, 2015). Trees forming ectomycorrhizae (ECM) with soil-borne fungi dominate temperate and boreal forest ecosystems (Brundrett, 2009). These mutualistic interactions rely on bidirectional exchanges of nutrients, which happen in mycorrhized roots. The host tree provides carbon derived from its photosynthesis to the fungus, whereas in return ECM fungi provide nitrogen, phosphorus and water. Ectomycorrhizal fungi thus inhabit a dual ecological niche, forest soils and tree root cells, requiring two contrasting ways of life: saprotrophic in soil (for nitrogen acquisition) and biotrophic within plant living tissues. Therefore, most ECM fungi develop (i) extramatrical mycelium exploring the rhizospheric surrounding soil, (ii) aggregated fungal hyphae ensheathing fine lateral roots named mantle, and finally (iii) a highly branched network of fungal hyphae (called the Hartig net) within the apoplastic space of epidermal and cortical root cells (Martin *et al.*, 2016). The Hartig net constitutes the biotrophic interface required for efficient nutrient exchanges. In particular environmental conditions (humidity, temperature), host-derived carbon can be used to build the fruiting body, a spore-releasing structure made of hyphal aggregation (Kües and Navarro-González, 2015; Genre and Bonfante, 2012; Lakkireddy *et al.*, 2011).

Despite their critical ecological roles, only a few ECM interactions have been studied at the molecular level. Notably, mechanisms mediating the early steps of ECM symbiosis development remain mostly uncharacterized (Daguerre *et al.*, 2017; Martin *et al.*, 2016). First, a pre-contact phase during which plant and fungi communicate is a prerequisite for successful root colonization. Diffusible molecules such as fungal auxins and sesquiterpenes likely mediate this communication and trigger an increase in the lateral roots formation (Felten *et al.*, 2009; Ditengou *et al.*, 2015; Krause *et al.*, 2015; Vayssières *et al.*, 2015). Then, fungal accommodation within the apoplast of root cells requires controlling both hyphal and host root development (Martin *et al.*, 2016). For example, in the poplar-*L. bicolor* model, both ethylene and jasmonic acid treatments restrain *in planta* fungal colonization i.e. restrict the intradical hyphal network (Plett *et al.*, 2014). The symbiotic interface at the Hartig net derives from remodeling of both fungal and plant cell walls (Balestrini and Kottke, 2016). Cell wall carbohydrates and proteins (e.g. hydrophobins, mannoproteins) are thus likely to actively contribute to *in planta* fungal colonization (for review, Balestrini and Kottke, 2016) and to efficient nutrient exchanges. Formation of the Hartig net also leads to a massive fungal colonization within the apoplast of colonized roots, without eliciting strong defence responses (Martin *et al.*, 2016). Considering ECM symbiosis as a biotrophic plant-fungal interaction, secreted fungal molecules likely govern plant colonization by subverting host immunity and manipulating its metabolism to promote the symbiosis establishment and/or functioning (Plett and Martin, 2015; Lo Presti *et al.*, 2015).

Genome-wide analysis of *Laccaria bicolor* has led to the identification of 98 proteins, named MiSSPs (Mycorrhiza-induced Small Secreted Proteins), up-regulated in symbiotic tissues (Martin *et al.*, 2008). Among them, only MiSSP7 has been described as a symbiosis effector so far. MiSSP7 is secreted by the fungus and it enters the host cells in which it localizes within the nucleus. Moreover, *MiSSP7*-RNAi mutants are impaired in ectomycorrhiza formation (Plett *et al.*, 2011). Inside the nucleus, MiSSP7 interacts with the *Populus trichocarpa* PtJAZ6 (JAsmonate Zim domain 6), a co-receptor of jasmonic acid (Plett *et al.*, 2014). MiSSP7 stabilizes PtJAZ6, avoiding its degradation in the presence of jasmonic acid, leading to the repression of target genes’ transcription (Plett *et al.*, 2014). Preliminary results have shown that genes involved in plant cell wall remodelling and plant defence responses might be these target genes (Plett *et al.*, 2014). Furthermore, effector proteins, such as SP7 from the arbuscular mycorrhizae fungus *Rhizophagus irregularis* (Kloppholz *et al.*, 2011), as well as PIIN_08944 and FGB1 (Fungal Glucan-Binding 1, PIIN_03211) from the root endophyte *Piriformospora indica* (Akum *et al.*, 2015; Wawra *et al.*, 2016) also suppress host immunity promoting root colonization and symbiosis. These data support the concept whereby a mutualistic symbiont uses its repertoire of secreted proteins to set up the symbiosis by suppressing host immunity and/or targeting cell-wall remodeling.

Considering the diversity of secreted proteins used by pathogens to alter the host metabolism, mycorrhizal fungi might use an equivalent strategy to setup their interaction. However, available literature on such proteins required for the symbiosis establishment is still very poor and functional analyses of MiSSPs are required to clarify and detail how mycorrhizal fungi communicate with their host plant to establish interaction (Plett and Martin, 2015). The Mycorrhiza-induced Small Secreted Protein of 8 kDa, i.e MiSSP8 (JGIv2 ID #388224), displays the third highest induction in mature ectomycorrhizal root tips and is also up-regulated in fruiting-bodies (Martin *et al.*, 2008). MiSSP8 is, with other MiSSPs, part of the “core” regulon expressed during the colonization of two hosts, *Populus trichocarpa* and *Pseudotsuga menziesii* (Plett *et al.*, 2015). The proteins associated to this core regulon have hence been hypothesized to be the key genetic determinants required for the symbiosis development in *L. bicolor* (Plett *et al.*, 2015). In this study, we report the functional analysis of *L. bicolor* MiSSP8 by using a combination of experimental and *in silico* approaches. Here, we identify MiSSP8 as a key symbiosis factor required for *L. bicolor* mycorrhization ability, mantle formation and subsequent Hartig net development. This symbiosis factor also has a repetitive motif at its C-terminus containing a kexin-like cleavage site, which is recognized by KEX2 *in vitro* and liberate four short peptides. The repetitive motif of MiSSP8 is found in other fungal proteins, mostly from ECM and saprotrophic fungi. All together our data indicate that MiSSP8 is involved in fungal mantle and Hartig net formation, potentially by regulating hyphal aggregation, and suggest a part of the symbiotic toolbox used by the ECM fungi to initiate symbiosis is already present in the genome of their saprotrophic ancestors.

## Material and methods

### Microorganism and plant material

*Saccharomyces cerevisiae* strains YTK12 (Jacobs *et al.*, 1997), MaV103 and MaV203 (Invitrogen) were propagated in YAPD medium (1% yeast extract, 2% peptone, 2% glucose, and 40 mg/L adenine) and cultured at 30°C. The ectomycorrhizal fungus *L. bicolor* Maire P. D. Orton strains (S238N and RNAi-lines) were maintained at 25°C on modified Pachlewski medium P5 +/− 150μg.ml^−1^ of hygromycin B (Pachlewski and Pachlewska, 1974, Di Battista *et al.*, 1996). The hybrid *Populus tremula x Populus alba* (INRA clone 717-1-B4) cuttings were micropropagated *in vitro* and grown on half MS medium (Murashige and Skoog, 1962) in glass culture tubes under a 16 h photoperiod at 24°C in a growth chamber. *L. bicolor* basidiocarps were harvested beneath inoculated Douglas fir in a nursery. Three fruiting body samples from two developmental stages were harvested; the “early” stage corresponding to just emerging fruiting body and “late” stage corresponds to ones with “open” cap. Samples from both developmental stages were divided into stipe and cape prior to RNAs extraction (Fig. S1).

To test for altered cell wall susceptibility, Congo red (150 μg/mL; Sigma-Aldrich) was added to the medium (Fig. S2).

### Yeast secretion trap assay

Functional validation of the predicted signal peptide of MiSSP8 was done using the yeast signal-sequence trap assay (Plett *et al.*, 2011). Briefly, full-length sequences of MiSSP8 with or without its signal peptide were cloned into pSUC2-GW, a plasmid carrying the invertase *SUC2* lacking both its initiation methionine and signal peptide. Yeast strain YTK12 was transformed with 200 ng of the plasmid using the lithium acetate method (Gietz and Schiestl, 2008). All transformants were confirmed by PCR with vector-specific primers and grown on yeast minimal medium with glucose (SD-W medium: 0.67% Yeast Nitrogen Base without amino acids, 0.075% tryptophan dropout supplement, 2% glucose and 2% agar). To assess invertase secretion, overnight yeast cultures were diluted to an O.D_600_ = 1 and 20 μl of dilution were plated onto YPSA medium (1% yeast extract, 2% peptone, 2% sucrose, and 1 μg.mL^−1^ antimycin A, inhibitor of cytochrome c oxidase). The YTK12 strains transformed with either the pSUC2-GW empty vector or containing SUC2SP (yeast invertase with signal peptide) were used as negative and positive controls, respectively.

### Genetic transformation of L. bicolor

The ihpRNA expression cassette/transformation vector was constructed using the pHg/SILBAγ vector system (Kemppainen *et al*, 2005; Kemppainen and Pardo, 2009). The full-lenght MiSSP8 cDNA was amplified from oligo(dT)18 synthesized S238N cDNA (First Strand cDNA Kit, (Fermentas) using the following gene specific primers:

MISSP8-*Sna*BIFor: CTTCTACGTAATGTATTTCCACACTCTTTTCG
MISSP8-*Hind*IIIRev: TGTCAAGCTTTCAATCACTATCGCGCCTC.

The cDNA was TA-cloned into pCR®2.1-TOPO® (Invitrogen) for sequencing and the corresponding plasmid was used as PCR template to obtain the amplicons needed for ihpRNA expression cassette construction. The cloning into the pSILBAy vector was carried out using the SnaBI, HindIII, BglII and StuI restriction sites in pSILBAy. Primers used for amplification of the MiSSP8 sequence arms were:

MISSP8-*Sna*BIFor: CTTCTACGTAATGTATTTCCACACTCTTTTCG
MISSP8-*Hind*IIIRev: TGTCAAGCTTTCAATCACTATCGCGCCTC
MISSP8-*Bgl*IIRev: TGTCAGATCTTCAATCACTATCGCGCCTC

Completed ihpRNA expression cassette was further cloned as a full length SacI linearized pSILBAy plasmid into the T-DNA of the binary vector pHg to create pHg/pSγMiSSP8. The pHg/pSγMiSSP8 was used for transforming *L. bicolor* dikaryotic strain S238N with *Agrobacterium tumefaciens* strain AGL1 (Kemppainen and Pardo, 2005). The transformed fungal strains were selected with 300 μg.mL^−1^ hygromycin B (Invitrogen) and were later maintained under 150 μg.mL^−1^ hygromycin B selection pressure on modified P5 medium. Four pHg/pSγMiSSP8 *L. bicolor* transformant strains were used for further molecular and physiological analyses.

### Molecular analyses of Laccaria bicolor transformants

Plasmid rescue of the right border (RB) - linked gDNA was carried out with BamHI cut and self-ligated *L. bicolor* gDNA according to Kemppainen *et al.* (2008). Sequencing of the rescued plasmids was done using M13/pUC-reverse primer (−26)17 mer. Left border (LB) TAIL-PCR was done according to Kemppainen *et al.*, 2009, using three T-DNA specific nested primers LB1.3 (Mullins *et al.*, 2001) and the arbitrary primer AD2 (Liu *et al.*, 1995). L3/AD2 amplified TAIL-PCR products were TA-subcloned into pCR®2.1-TOPO® (Invitrogen) and sequenced with the L3 primer. All the PCR reactions were carried out using Tpersonal thermocycler (Biometra ®) and PCR chemicals from Fermentas. Sequencing reactions were purchased from Macrogen Sequencing Service (Seoul, SK).

### In vitro mycorrhization experiments

For *in vitro* mycorrhization tests between *P. tremula x alba* INRA717-1B4 and *L. bicolor*, we used a “sandwich” co-culture system described in Felten *et al.* (2009). After three weeks of co-incubation of poplar cuttings with *L. bicolor*, at least 10 to 20 biological replicates were analysed for the percentage of colonized roots with the *L. bicolor* wild-type strain S238N or with the four *L. bicolor MiSSP8*-RNAi lines. Two independent empty vector transformants of *L. bicolor* (ev7 and ev9 lines, Plett *et al*, 2011) were also tested for their ability to colonize roots.

### Motif analysis

For the identification of proteins sharing the DWRR motif found in MiSSP8 sequence, the following regular expression DW[K/R]-x(2,20)-DW[K/R]-x(2,20)-DW[K/R]-x(2,20)-DW[K/R]-x(2,20) was used on a set (Table S1) of fungal and oomycete proteomes available at the MycoCosm database (March 2019) using the PS-Scan software (de Castro *et al*, 2006) with the options -g (greedyness off) -v (overlaps off) -p (pattern search). Only protein sequences from published genomes starting with a methionine were kept for further analysis (Table S1). In order to assess conservation of the DWRR motif, protein sequences retrieved by PS-Scan were further analyzed by GLAM2 (Gapped Local Alignments Motifs) software v 4.11.0 (Frith *et al.*, 2008) with default parameters. Similar GLAM2 analysis was performed on shuffled sequences as control. The presence of a signal peptide in the protein sequence was assessed with SignalP v 4.1 (Petersen *et al*, 2011) with default parameters.

### Microscopy analysis of poplar-ECM roots

ECM root tips of *poplar-L. bicolor* (WT or *MiSSP8*-RNAi lines) were fixed 24 h in 4% paraformaldehyde in PBS buffer (100 mM phosphate buffer, 2.7 mM KCl and 137 mM NaCl pH 7.4) at 4°C. The root segments were embedded in agarose 5% and cut into 25 μm radial sections with a Leica VT1200S Leica vibratome (Leica Microsystems). Sections were categorized according to their distance (100, 200 and 600 μm) from the root apex. 25 μm-width sections were stained with 10μg.mL^−1^ wheat germ agglutinin (WGA)–Alexa Fluor® 488 Conjugate (W21404, ThermoFisher, France) and 1μg.mL^−1^ propidium iodide (Invitrogen, France). To compare the development of the Hartig net between samples, sections between 200 and 600 μm distance from the root apex were analyzed.

### Cell imaging by confocal laser-scanning microscopy

Transversal sections of ECM root tips were viewed by a Zeiss LSM780 (Carl Zeiss AG, Germany) confocal laser scanning microscope system. Images were obtained with objective CAPO-40x/1.2 water-immersion objective, acquired sequentially to exclude excitation and emission crosstalk (when required). Spectral deconvolution was used to assess specificity of the emission signal. Images were processed with ZEN (Carl Zeiss AG) software.

### Quantitative RT-PCR

For the expression of *MiSSP8* in colonized root tips, extramatrical and free-living mycelium, total RNA was extracted from 100 mg biological material using the RNeasy Plant Mini Kit (Qiagen), extraction buffer RLC was supplemented with 2% PEG8000. An on column DNase I treatment was included in the protocol. 500 ng of total RNA was converted into cDNA in a 20 μl reaction using the High-Capacity cDNA Reverse Transcription Kit (Applied Biosystems, Life technologies, Thermo Fisher Scientific) according to manufacturer's instructions and subsequently 1/5 diluted in sterile RNAse and DNAse free water. Real-time qPCR was performed in an optical 96-wells plate with a StepOne sequence detection system (Applied Biosystems, Life Technologies) and fast cycling conditions (20 s at 95°C, 40 cycles of 3 s at 95°C and 30s at 60°C). Each 10 μl reaction contained 2X Fast SYBR green Master Mix (Applied Biosystems, Life Technologies), 300 nM gene-specific forward and reverse primers, water and 10 ng of cDNA. Data were expressed relatively to the sample with the highest expression level (2 ^−(Ct-Ctmin)^) and normalized against six reference genes (JGIv2 IDs #293350, #611151, #313997, #446085, #246915 and #319764). Stability of the reference genes was confirmed by geNorm analysis (Vandesompele *et al.*, 2002). The normalization factor NF for each sample was calculated as the geometric mean of the relative expression level of the six reference genes. Consequently, the expression level of *MiSSP8* for an individual sample was obtained by the formula 2 ^−(Ct-Ctmin)^/NF according to Vandesompele *et al.* (2002). Finally, data were rescaled relative to the FLM.

For the fruiting body expression, cDNA was obtained from 500 ng of total RNA using the i-Script cDNA reverse transcription kit (Biorad) in a final volume of 20 μL. RT-qPCR reactions were performed on 10 ng cDNA and 300 nM forward and reverse primers in each reaction, using the RotorGene (Qiagen) with the standard cycle conditions: 95 °C for 3 min; 40 cycles at 95 °C for 15 s and 65 °C for 30 s, followed by a melting curve analysis (temperature range from 65 °C to 95 °C with 0.5 °C increase every 10 s). In this particular case, transcript abundance was normalized using *L. bicolor* histone H4 (JGIv2 ID# 319764) and ubiquitin (JGIv2 ID #446085) encoding genes. Amplification efficiency (E) was experimentally measured for each primer pair and was taken in account for calculation of normalized expression (Pfaffl *et al.*, 2001).

### Production of recombinant protein and biochemical analysis

*MiSSP8Δ1*-20 (i.e. devoid of its first twenty amino-acid residues) was synthesized by Genecust (Luxembourg) and subcloned into the pET-28a-CPDSalI vector (Shen *et al.*, 2009) between *Nco*I and *Sal*I restriction sites. The resulting plasmid was subsequently used for the transformation of the Rosetta2 (DE3) pLysS strain of *E. coli* (Novagen). The expression of the recombinant protein (ending with DSDVD in C-ter) was performed at 37°C in Lysogeny broth (LB) medium supplemented with 30 μg.mL^−1^ of kanamycin and 34 μg.mL^−1^ of chloramphenicol. When the cell culture reached an O.D_600_ of 0.7, recombinant protein expression was induced by the addition of 0.1 mM isopropyl β-D-1-thiogalactopyranoside (IPTG), and the cells were grown for further 4 hours at 37°C. Cells were then harvested by centrifugation (13000rpm, 5min), resuspended in a 30 mM Tris/HCl pH 8.0, 200 mM NaCl lysis buffer and lysed by sonication. The cell extract was centrifuged at 35000 g for 25 min at 4°C to remove cellular debris and aggregated proteins. C-terminal His-tagged proteins were purified by gravity-flow chromatography on a nickel nitrilotriacetate (Ni-NTA) agarose resin (Qiagen) according to the manufacturer’s recommendations followed by an exclusion chromatography on a Superdex75 column connected to an ÄKTA Purifier™ (GE Healthcare). CPD-tag was cleaved using 200 μM inositol-6-phosphate as described by Schen *et al.* (2009) and removed by size exclusion chromatography. Circular dichroism experiments were carried out with 75 μM of recombinant MiSSP8 in 10 mM sodium phosphate buffer pH 7 using a Chirascan Plus (Applied Photophysics).

### *In vitro* digest assay of recombinant MiSSP8 by KEX2

Recombinant MiSSP8 protein (75μg) was incubated with 0.02 units of recombinant KEX2 protease (MoBiTec) in 5 mM CaCl_2_, 50 mM Tris pH7 at room temperature during 1, 2, 4 or 6 hours then was stopped by heating at 95°C for 10 minutes. 50% of each samples was analysed on Tris-Tricine-precast 12-15% polyacrylamide gels (Biorad), stained with Coomassie blue.

### Mass spectrometry analysis using LC Q-TRAP of KEX2-digested MiSSP8

Synthesized peptide standard (DSDW) obtained from Genecust was solubilized and diluted to 10^−8^M in 2% acetonitrile in water and stored at −20°C. *In vitro* KEX2/MiSSP8 digest assay was diluted in 100 μl of 2% acetonitrile in water and stored at −20°C. The U-HPLC 3000 (Dionex) was equipped with a reverse-phase column Acquity UPLC BEH-C18 (2.1 × 150 mm, 1.7 μm, Waters). Separation was done with a gradient of eluent A (water/0.1% formic acid) and eluent B (acetonitrile), started at 5% B for 1 min, followed by a 8 min gradient to 100% B, followed by an isocratic step at 100% B for 2 min, a 2 min gradient back to 5% B, equilibration step 2 min at 5% B, before starting another analysis, at a constant flow rate of 300μl/min. 10μl of each samples were injected.

The mass spectrometer used was a 4500 Q-Trap mass spectrometer (Applied Biosystems, Foster City, USA) with an electro-spray ionization source in the positive ion mode. The capillary voltage was fixed at 4500 V and the source temperature at 400°C. Optimizations of the source parameters were done using the peptide standards DSDW at 10^−5^ M in 2 % acetonitrile by infusion at 7μl. min-1, using a syringe pump. For MS/MS analysis, enhanced product ions (EPI) mode was used. Declustering potential was fixed at 110 V. Fragmentations were induced by collision induced dissociation (CID) with nitrogen at a collision energy of 40 V. For the multiple reaction monitoring (MRM) mode, each precursor ions and b/y ions were calculated for the predicted peptide using Protein Prospector. For relative quantification, the transitions selected for each peptide were: 522.5>185 (DSDW, RT: 5.0 min), 550.5>175 (DSDVD, RT: 4.5 min), 678.5>254 (RDSDW, RT: 5.0 min), 678.5>361 (DSDWR, RT: 4.4 min), 834.5>517 (DSDWRR, RT: 4.0 min), 834.5>361 (RDSDWR, RT: 4.0 min), 834.5>313 (RRDSDW, RT: 4.0 min). The intensity of peak height was measured in counts per second (cps).

## Results

### *MISSP8* is up-regulated both in ECM root tips and fruiting body

Transcript profiling of *L. bicolor* free-living mycelium and *P. trichocarpa* colonized root tips have highlighted the presence of >50 Mycorrhiza-induced Small Secreted Proteins (MiSSPs) in this ECM fungus. Of these, *MiSSP8* displayed the third highest induction in mature ectomycorrhizal root tips (Martin *et al.*, 2008). To determine the regulation of MiSSP8 throughout ECM development, we investigated *MiSSP8* expression during *in vitro* ECM time course with *P. tremula x alba* using real time-qPCR. At 7 days post-contact, fungal hyphae started colonization of fine roots and form the mantle. At 14 days, the Hartig net is present and at 21 days, fully mature ECM tissues have developed. The very low and constitutive-level of *MiSSP8* expression in free-living mycelium (FLM) was set as a reference. *MiSSP8* was up-regulated in ECM root tips from day 7 to day 14, reaching its maximum induction at this latter time point (Fig. 1A). The expression decreased to reach the same level as in FLM at 21 days (mature ECM). Expression of *MiSSP8* in the extraradical mycelium (i.e. the part of the rhizospheric mycelium not in contact with the roots) was the same as FLM all along the time course (Fig. 1A). In order to investigate the expression level of *MiSSP8* in a non-symbiotic tissue, we performed qRT-PCR on *L. bicolor* fruiting body tissues (stipe and cap) at two different developmental stages (early and late) (Fig. 1B, Fig. S1). The expression of *MiSSP8* in *L. bicolor* sporocarps is higher than its expression in FLM and ECM root tips. Overall, *MiSSP8* was strongly induced during fruiting body-development and *P. tremula x alba* root colonization (mantle and Hartig net formation).

**Figure 1:**
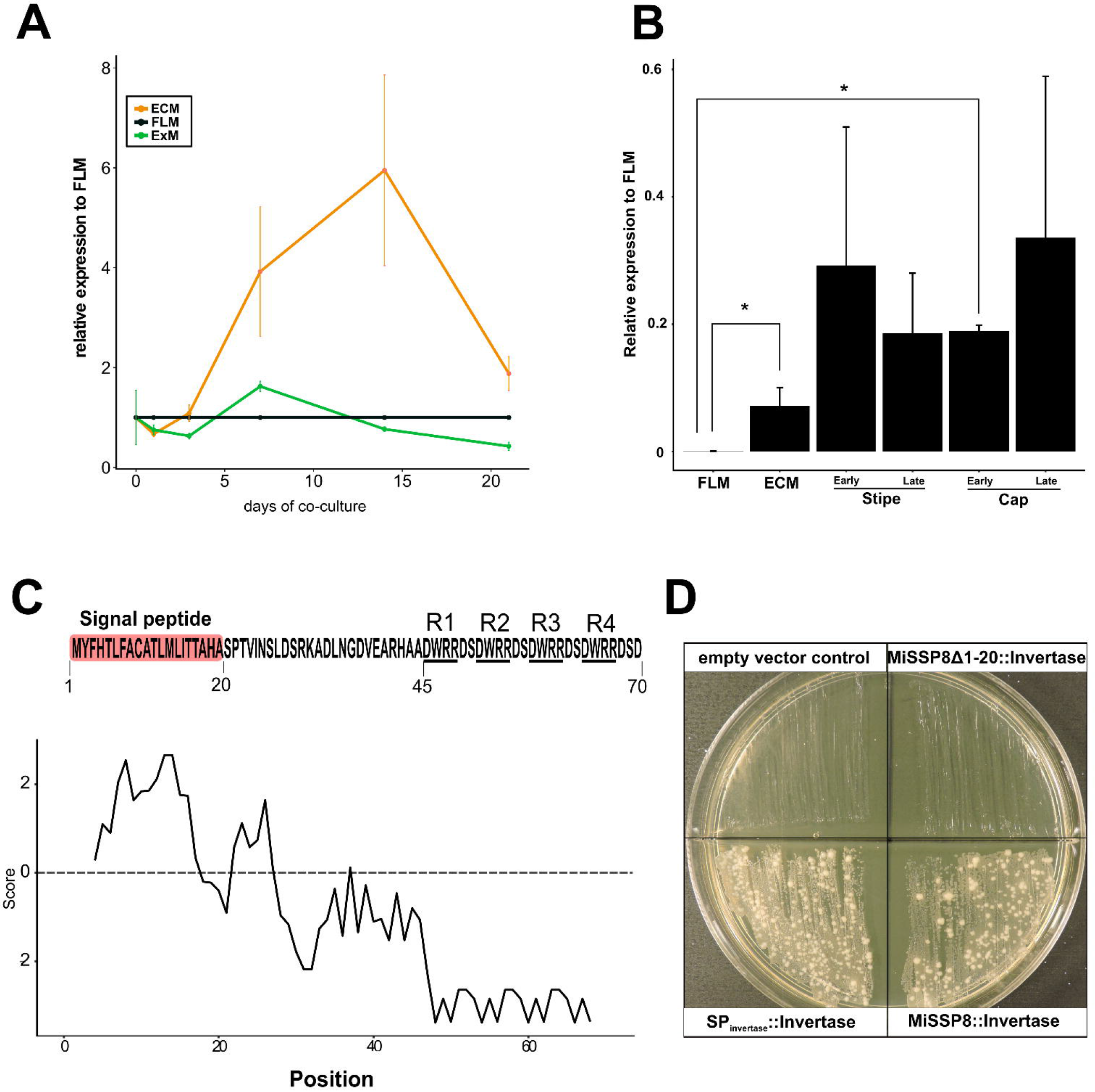
MiSSP8 is a secreted repeat-containing protein highly expressed in ectomycorrhizal root tips and fruiting body tissues. **(A)** Real-time qPCR time course performed on *Laccaria bicolor* extraradical mycelium (ExM), free-living mycelium (FLM) and during development of *Populus tremula* x *alba* ECM root tips. Expression of *MiSSP8* in FLM was set as a reference. Error bars represent standard deviation from three biological replicates. **(B)** Real-time qPCR analysis on the expression of *MiSSP8* versus two reference genes in the fruiting body of *L. bicolor*, the cap and stipe, at both early and late stages of development. Stars indicate significant differences (p-value cut-off < 0.05) using Welch t-test (Welch, 1947). Error bars represent standard deviation from three biological replicates. **(C)** MiSSP8 sequence, domain organization and hydrophobicity score as obtained by Protscale using the amino-acid scale from Kyle and Doolittle (1982) with a window size of 5. **(D)** Yeast signal trap assay shows that the signal peptide predicted in MiSSP8 sequence is functional in yeast. *Saccharomyces cerevisiae* was transformed either with empty vector control (top left), mature MiSSP8 (lacking its signal peptide) fused to mature invertase (MiSSP8Δ1-20::Invertase, top right), full-length invertase (with its signal peptide, bottom left) or full length MiSSP8 fused to mature invertase (MiSSP8::Invertase, bottom right). Transformed yeasts were plated on growth medium containing sucrose as the sole source of carbon.

### MiSSP8 is secreted as indicated by the yeast invertase secretion assay

MiSSP8 is a protein containing 70 amino acids, the first twenty residues of which encode a predicted signal peptide as predicted by SignalP v4.1 (Fig. 1C). According to the hydrophobicity plot, the mature MiSSP8 is a hydrophilic protein with charged amino acids exposed to solvent (Fig. 1C) but there is no predicted secondary structure (Fig. S2A). A circular dichroism experiment performed on the recombinant protein produced in *E. coli* further confirms the lack of secondary structure of MiSSP8 (Fig. S2B). In order to confirm that the predicted signal peptide is functional and properly processed, the full length MiSSP8 (including its signal peptide) was fused to the yeast invertase *SUC2*, which catalyzes the hydrolysis of sucrose. The transformed yeasts were able to grow on a minimal medium supplemented with sucrose and antimycin (Fig. 1D, bottom right) like the yeast transformed with a full-length invertase (Fig. 1D, bottom left). Yeasts transformed with an empty vector control (Fig. 1D, top left) or an invertase fused to MiSSP8 lacking its signal peptide (Fig. 1D, top right) did not grow on the same medium. This demonstrates that the signal peptide of MiSSP8 is properly recognized and processed in yeast and triggered the secretion of the invertase. Therefore, it is likely that *L. bicolor* secretes MiSSP8 into the extracellular space during root colonization and fruiting body development.

### MiSSP8-repetitive motif is shared with proteins containing repeats from saprotrophic and ectomycorrhizal fungi

The mature MiSSP8 is composed of 50 amino acids, with a predicted molecular weight of 6105.32 Da and a theoretical pI of 5.12. At its C-terminus, MiSSP8 contains a 25 amino acids long sequence carrying a repetitive motif (i.e. DWRR), repeated four times consecutively. This protein displayed sequence similarities with only one protein of *Laccaria amethystina* (JGIv2 #676588), a close relative of *L. bicolor*, according to BLASTP search on NCBI and JGI MycoCosm databases. Altogether, these data suggest that MiSSP8 is a natively unstructured protein without sequence similarities with previously characterized proteins. To identify additional proteins with a similar motif at their C-termini, we used a pattern search algorithm and identified in total 38 proteins from 23 published fungal proteomes (Fig. 2, Table S1). The DW[K/R]R containing proteins identified were mostly associated to saprotrophs (32/38) and ectomycorrhizal (5/38) fungi including also one arbuscular mycorrhizal fungus, but none pathogenic fungi (Fig. 2, Table S1). Most of the proteins identified above using pattern search algorithm (34/38) had a predicted signal peptide (Fig. 2). No protein domains were detected by PROSITE database, except for two proteins from the white rot fungus *Schizophyllum commune*, which possess a N-terminal aspartic peptidase A1 domain (Fig. 2). GLAM2 motif analysis found an enrichment of a DWR/KR motif, named DW[K/R]R thereafter.

**Figure 2:**
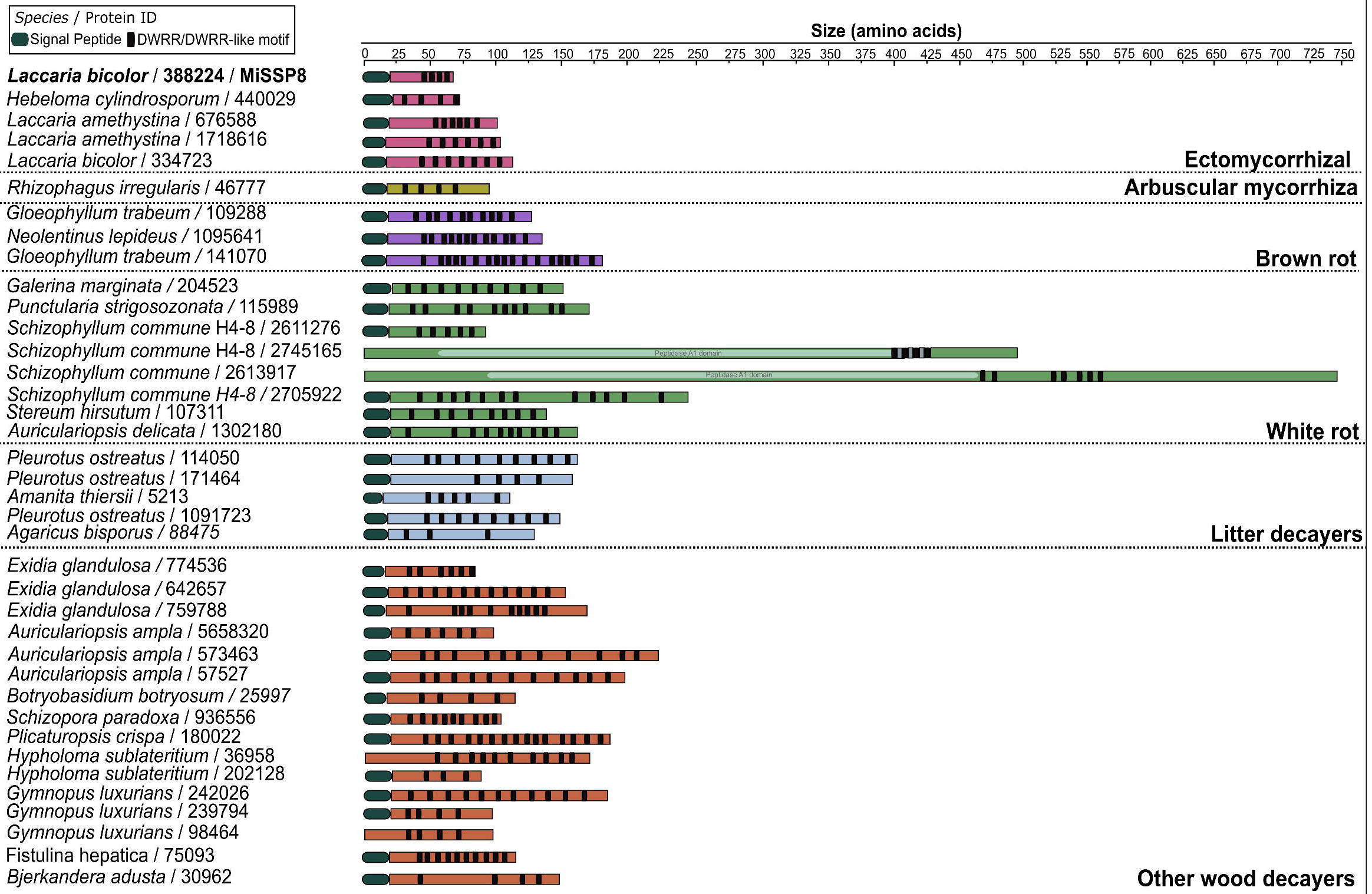
The DW[K/R]R proteins possess a repetitive structure and are likely secreted. The modular structure of the proteins is shown. Protein domains have been identified by PROSITE and signal peptides are predicted by SignalP v4.1.

The median size of the proteins was 138.5 amino acids, ranging from 70 to 738 amino acids (Fig. 3A). The number of repetitions of the conserved peptide varied from three to seventeen, with no correlation between the size of the protein and the number of motifs (Fig. 3B, Fig. S3). RNA-Seq expression data from *Pinus pinaster* root tips colonized by the ectomycorrhizal fungus *Hebeloma cylindrosporum* showed that the DWRR containing protein from *H. cylindrosporum* (JGI IDv2: 440029) was upregulated during root colonization (Doré *et al.*, 2015; GEO accession number GSE63868/GSE66156). GLAM2 motif analysis found an enrichment of a DWR/KR motif, named DW[K/R]R thereafter (Fig. 3C, Fig. S3).The DW[K/R]R repetitive motif is therefore shared between proteins predicted as secreted for the most part and associated to saprotrophs and mycorrhizal fungi.

**Figure 3:**
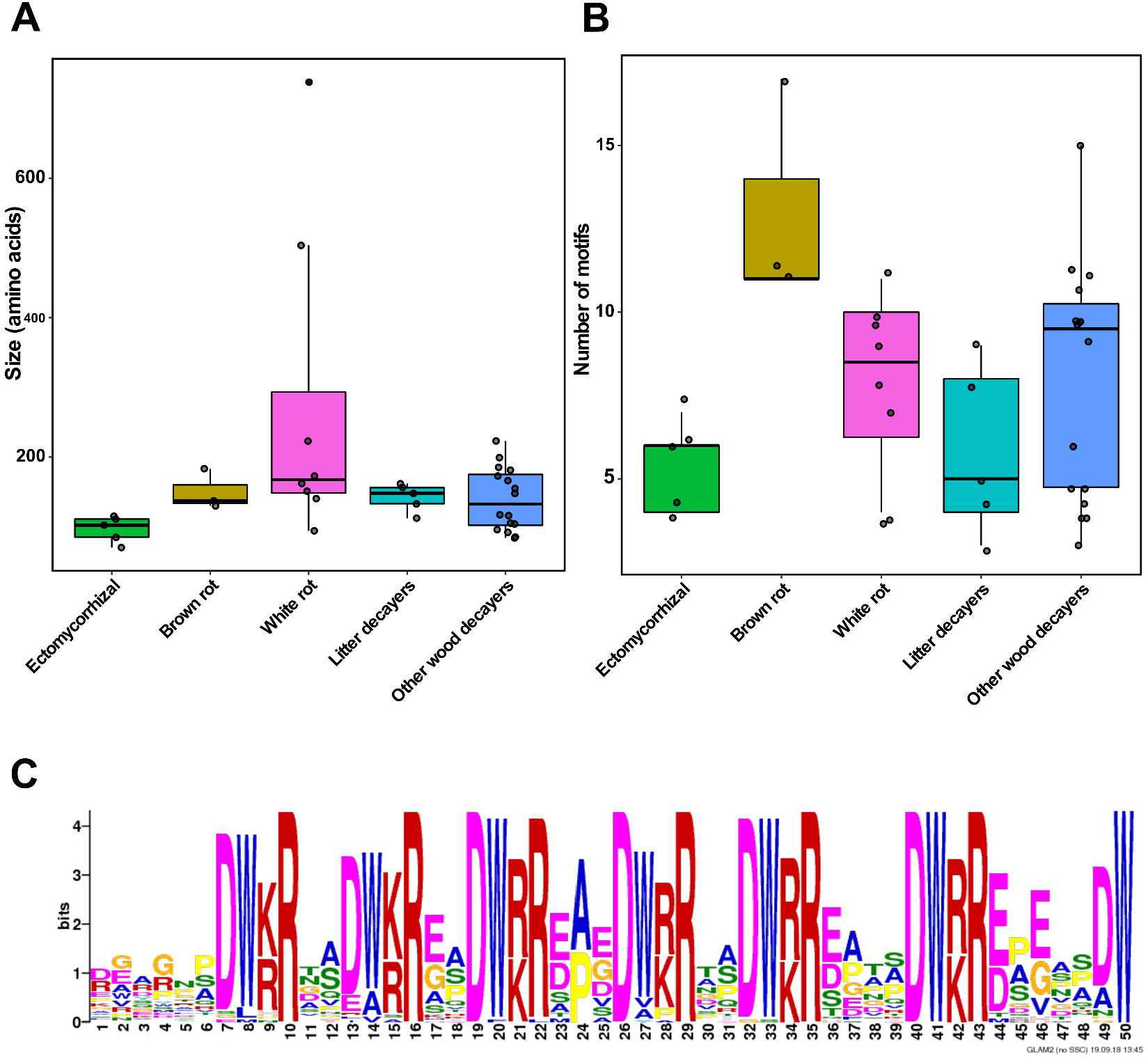
MiSSP8-repetitive motif is found in proteins from saprotrophic and ectomycorrhizal fungi. **(A)** Size distribution (in amino acids) of the DW[K/R]R-containing proteins within the different fungal lifestyles identified. **(B)** Number of repetitions identified in the DW[K/R]R-containing proteins. **(C)** Identification of the DWRR motif by GLAM2 software. The set of 38 sequences retrieved with PS-Scan has been submitted to GLAM2 run with default parameters.

### MiSSP8 possess a repetitive motif containing a kexin cleavage site, recognized *in vitro* by the yeast KEX2 protease

The presence of a repetitive motif in the protein sequence suggests that MiSSP8 might be processed post-translationally in order to become active. Interestingly, the motif DW[K/R]R contains [K/R]R residues, which are known recognition sites for KEX2 proteases in fungi (Mizuno *et al.*, 1988; Mizuno *et al.*, 1989). In fungi, an additional proteolytic cleavage of the KEX2-peptides occurs by KEX1 to withdraw the two amino acids KR or RR. The processing of MiSSP8 at the identified cleavage sites (i.e. RR at each DWRR repeats) by KEX1 and KEX2 would release one peptide of 27 amino acids, three short peptides of four amino acids (DSDW) and a final three amino acids long peptide DSD. The sole action of the KEX2 protease will lead to the release of the DSDWRR peptide. In order to assess whether this motif is recognized by KEX2, we performed *in vitro* digest assay of recombinant mature MiSSP8 with recombinant yeast KEX2 protein. MiSSP7, a *L. bicolor* small-secreted protein of 7kDa, which does not contain the (K/R)R motif, was used as negative control. After 2h of incubation, the amount of undigested MiSSP8 decreased and was not detectable after 4h of incubation. For the same incubation period, MiSSP7 was not degraded (Fig. 4). Using mass spectrometry, we assessed whether the predicted DSDWRR or additional peptides were produced to confirm the specificity of the *in vitro* enzyme test. The separation and analytical detection parameters by mass spectrometry were fixed using the synthetic standard DSDW. We obtained a limit of detection of the standard DSDW at 10^−8^ M for 10 μl injected, at a retention time of 5.08 min. In the products of digestion, we searched for C-terminal DSDVD, the peptide DSDW and other predicted peptides with one to two additional arginine (R) before/after DSDW (Figure 1C). Among the major detected peptides, we observed the C terminal peptide DSDVD, DSDWR and DSDWRR, meaning that as predicted, KEX2 can hydrolyze MiSSP8 after two arginines (formation of DSDVD and DSDWRR), and between the two remaining arginines, to form DSDWR (Fig. S4).

**Figure 4.**
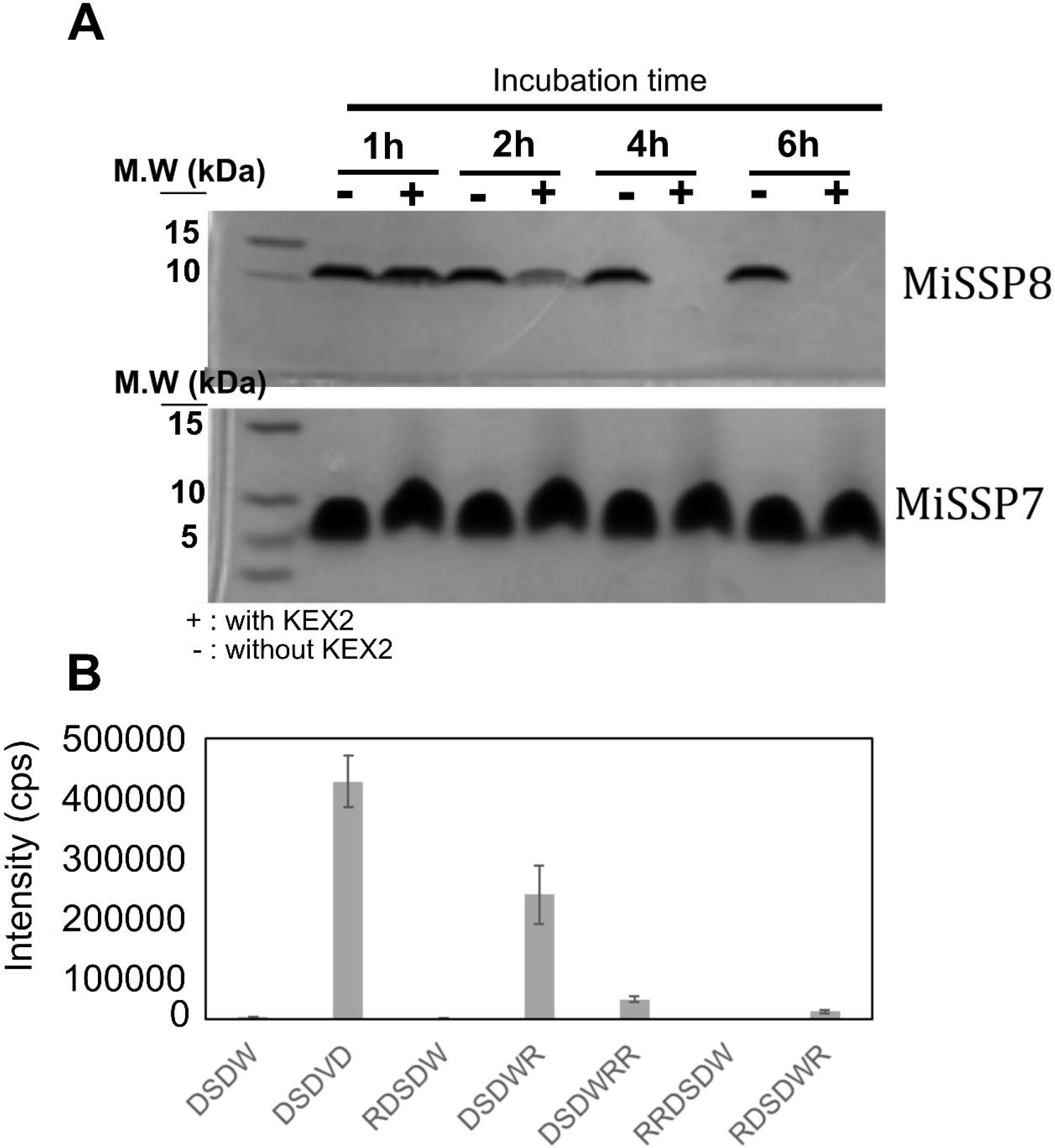
Recombinant MiSSP8 is cleaved by KEX2 protease *in vitro*. **(A)** Visualization of proteolytic digests of recombinant MiSSP8 and MiSSP7 proteins by recombinant KEX2 protease, separated on Coomassie-stained-polyacrylamide gels. A time course over 1, 2, 4 and 6h of incubation was performed. Sizes are indicated in kDa. MiSSP7 is used as negative control since it is not a putative substrate for KEX2 protease.+ and − indicate the presence or absence of KEX2, respectively. **(B)** Detected peptides by LC-MS/MS after 4h of digestion by the recombinant KEX2 protease of the MiSSP8 protein. Theoretical formed peptides are DSDWRR and DSDVD but additional combinations of cleavages were hypothesized. The transitions selected for each peptide were: *m/z* 522.5>185 (DSDW, RT: 5.0 min), 550.5>175 (DSDVD, RT: 4.5 min), 678.5>254 (RDSDW, RT: 5.0 min), 678.5>361 (DSDWR, RT: 4.4 min), 834.5>517 (DSDWRR, RT: 4.0 min), 834.5>361 (RDSDWR, RT: 4.0 min), 834.5>313 (RRDSDW, RT: 4.0 min). Peak height intensity is measured in counts per second (cps). Mean of 3 independent replicates, standard error of the mean is reported.

### RNAi-mediated knockdown of MiSSP8 encoding gene impairs mycorrhization rate

Since *MiSSP8* expression is induced during the early steps of ectomycorrhizal symbiosis development, we assessed whether MiSSP8 is required for establishment of symbiotic interaction. Generation of knockout mutants through homologous recombination has not been accomplished so far in *L. bicolor*. However, genetic tools to obtain *L. bicolor* strains with significantly reduced target gene expression levels through RNA interference (RNAi) are available (Kemppainen *et al.*, 2009; Kemppainen and Pardo 2010). RNAi hairpin targeting the MiSSP8 transcripts was expressed in *L. bicolor* through *A. tumefaciens* mediated transformation (ATMT). A transgenic ATMT *Laccaria* library was generated and 24 randomly selected independent RNAi lines were passed through consecutive hygromycin B selection steps. Among these, four *Laccaria* RNAi lines were analyzed at molecular level (Plett *et al*, 2011; Table S2). Real-time qPCR analysis confirmed down-regulation of *MiSSP8* in the four transgenic lines, with a reduction from 82% to 95% compared to the empty-vector control lines (Fig. 5A). We quantified the ability of these transgenic RNAi lines to colonize *P. tremula x alba* roots *in vitro*. The mycorrhization rate, defined as percentage of ECM root tips formed over the total number of lateral roots, was found significantly reduced with the mutants, being 45% for the wild-type and empty-vector controls and only 10 to 15% for the four *L. bicolor MiSSP8*-RNAi lines (Fig. 5B). The mycelial growth of each of the *MiSSP8*-RNAi line was tested on rich agar medium or on Congo Red-containing medium and it was similar to the wild-type strain S238N (Fig. S5). Neither did the RNAi mutant show any growth defect nor altered fungal cell wall susceptibility compared to the wild-type fungus. Altogether, it demonstrates MiSSP8 is required for the establishment of the symbiosis with poplar.

**Figure 5:**
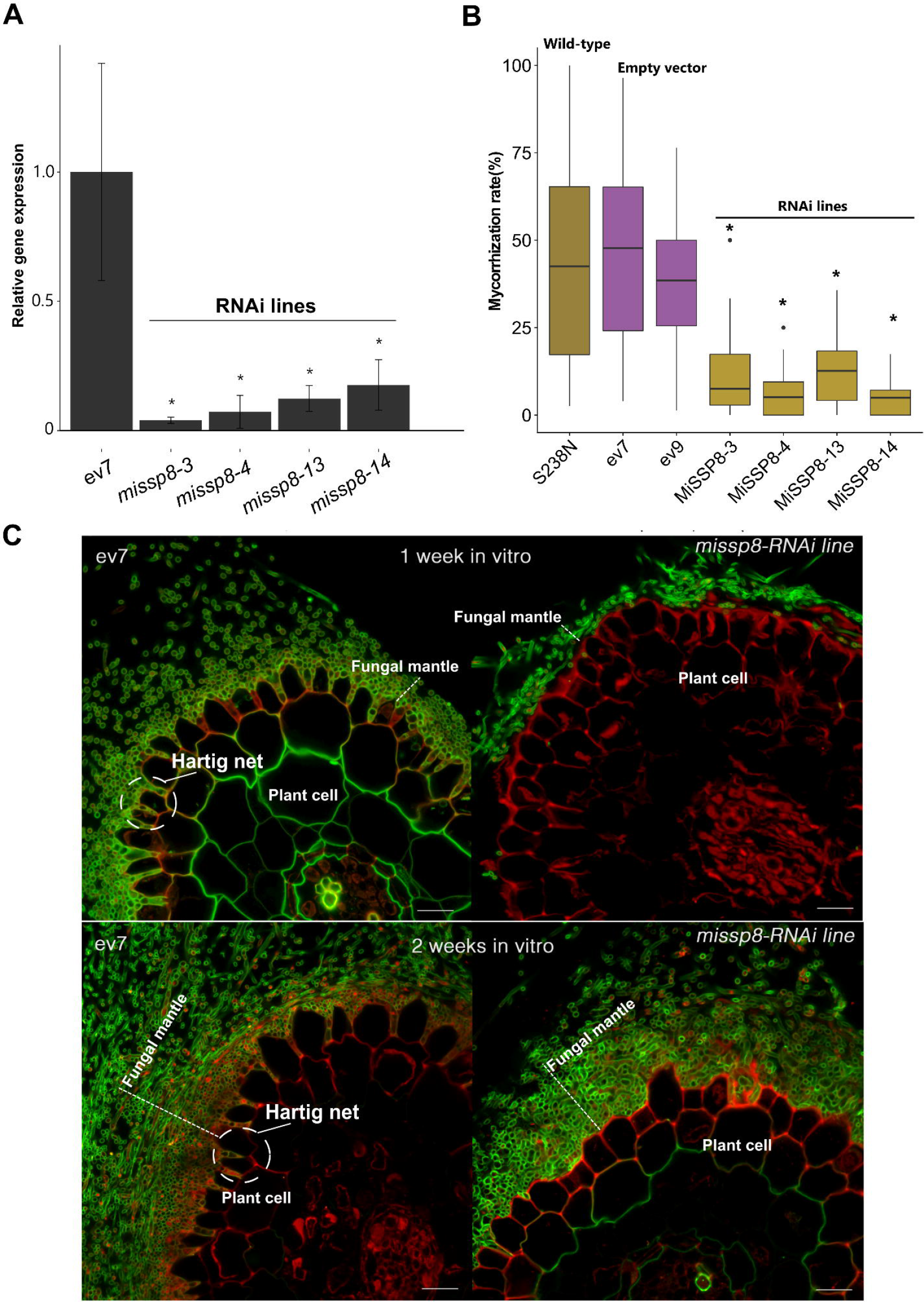
Knockdown of *MiSSP8* impairs the establishment of ectomycorrhizal symbiosis. **(A)** Expression was measured by qRT-PCR. All values are shown as mean ± standard deviation; n = 3. Stars indicate that expression is statistically different than control line based on a non-parametric Kruskal-Wallis test followed by a Games-Howell post-hoc test (Ruxton and Beauchamp, 2008). Adjusted p-value (FDR) cut-off = 0.05. ev indicates fungal lines transformed with an empty vector (i.e. negative control). **(B)** Knockdown of *MiSSP8* strongly decreases the number of ectomycorrhizal root tips compared to the wild-type strain or the empty vector controls. Percentage of poplar (*Populus tremula x alba*) mycorrhizal root tips formed by wild-type *L. bicolor* S238N and two empty vector transformation controls (ev7 and 9) versus four independent *L. bicolor MiSSP8*-RNAi lines. All values are shown as mean ± standard deviation; n=25; Stars indicate significant difference from wild-type, ev7 and ev9 using t-test and a p-value cut-off < 0.05. **(C)** Transversal section of ectomycorrhizal root tips formed by ev7 control transformant line and *MiSSP8-*RNAi line 3. The *MiSSP8-*RNAi line 3 displays a loose mantle and no Hartig net compared to the empty vector control strain (ev7). Scale bars: 10 μm. Propidium iodide stains plant cell walls and nuclei (red) and WGA-Alexa 488 stains fungal cell walls (green). Green signal in root cortical cells correspond to aspecific WGA-Alexa 488 fixation to plant cell walls and not to fungal cells.

### RNAi-mediated knockdown of *MiSSP8* impairs formation of fungal mantle and subsequent Hartig net development

To estimate whether MiSSP8 is necessary for fungal mantle and Hartig net development (two hallmarks of the mature ectomycorrhizal root tips), we performed microscopy analysis on ECM poplar root tips formed. After one week of contact with poplar roots, the fungal mantle formed by *MiSSP8*-RNAi lines was strongly unstructured and thinner than the one obtained with the reference strain (empty vector (ev) control). In addition, whereas ev control already formed first steps of Hartig net, the *MiSSP8*-RNAi lines did not display any kind of fungal colonization. After two weeks, *MiSSP8*-RNAi lines displayed one or two layers of fungal hyphae forming the mantle (Fig. 5C). This observation contrasts with the fungal mantle formed with the control transformant strain, which displayed a thick and well-organized fungal sheath with several layers of fungal hyphae stacked on each other. More strikingly, *MiSSP8*-RNAi lines were impaired in their ability to form the Hartig net even after two weeks of contact (Fig. 5C). Altogether, these results highlight the involvement of MiSSP8 in the differentiation of the fungal mantle precluding hyphal expansion to form the Hartig net.

## Discussion

### MiSSP8 is required for symbiosis development, likely through its role for hyphal aggregation

Several MiSSPs from *L. bicolor*, including MISSP8, are known to be part of the “core” regulon expressed during the colonization of two hosts, *Populus trichocarpa* and *Pseudotsuga menziesii* (Plett *et al.*, 2015). These proteins have hence been hypothesized to be key genes required for the symbiosis development in *L. bicolor* (Plett *et al.*, 2015). The strong decrease in the ability to form mycorrhizae by the *MiSSP8*-targeted RNAi lines is consistent with this hypothesis. In addition, our study clearly shows that MiSSP8 is directly involved in the symbiosis establishment, through a role played in mantle formation and the subsequent Hartig net development.

Real-time qPCR performed on free-living mycelium, symbiotic tissues and fruiting bodies showed a high level of *MiSSP8* expression both in mycorrhizal root tips and fruiting bodies but not the free-living mycelium. This expression profile suggests an involvement of MiSSP8 in both symbiosis-related (i.e. formation of ectomycorrhiza) and non-symbiosis-related processes (i.e. fruiting body formation). Both the ectomycorrhizal mantle sheath and the fruiting body tissues are composed of pseudoparenchyma, a pseudo-tissue made of aggregated hyphae that looks like the plant parenchyma (Brunner and Scheidegger, 1992; Peterson and Farquhar, 1994). Ectomycorrhizae developed by *MiSSP8-*RNAi lines display a disorganized fungal mantle and no Hartig net formation. We speculate that this phenotype results of the lack of fungal aggregation. Hyphae from mantle are indeed glued together and embedded into an extracellular material composed of glycoproteins and fungal polysaccharides (e.g. chitin, β 1-3 glucans) (Massicotte *et al.*, 1990; Dexheimer *et al.*, 1994). Previous studies propose a sequential role of hydrophobins, repellent and polypeptides with a RGD motif (e.g. SRAP32), in the aggregation of hyphae during mantle development in the ectomycorrhizal fungus *Pisolithus tinctorius* (reviewed by Martin *et al.*, 1999).

### MiSSP8 is repetitive protein with a kexin cleavage site sharing similarities with other repetitive proteins from saprotrophs

Since *MiSSP8* displays high sequence similarity with only one *L. amethystina* gene, we conclude that MiSSP8 is a *Laccaria*-specific gene, upregulated both in symbiosis and in fruiting bodies. These lineage-specific genes may have been formed *de novo* or may derived from neofunctionalization of duplicated genes or from ancestral genes that have strongly diverged due to selection pressure (Kohler *et al.*, 2015; Pellegrin *et al*, 2015). However, despite the absence of sequence similarities, a MCL (Markov Cluster Algorithm) analysis performed on a set of 49 fungal genomes had previously retrieved 33 proteins containing a similar motif as MiSSP8 (Kohler *et al*, 2015). With the current analysis, and using a different search approach. we also identify additional fungal proteins harboring the same repetitive motif than MiSSP8 as well as a kexin endoproteinase cleavage site (K/R)R (Mizuno *et al.*, 1988; Mizuno *et al.*, 1989). In addition, we showed this cleavage site is recognized *in vitro* by yeast KEX2. Despite several trials, we were not able to detect the predicted released peptides in ectomycorrhiza root tips or *L. bicolor* fruiting bodies (data not shown). This could be due to post-translational modifications of the peptides or a fixation to extracellular components such as fungal cell wall carbohydrates or glycoproteins. We can therefore only suggest that the proteins containing the (DWRR)_n_ motif could be processed by kexin prior to their secretion or go through post-translational modifications in order to become active and release such peptides. Several fungal peptides are produced from KEX2-processed precursor proteins (Le Marquer *et al*, 2019) e.g., cyclic peptides with mycotoxic activity in Ascomycota such as phomopsins (Ding *et al.*, 2016) or ustiloxins (Umemura *et al.*, 2014). On the other hand, *Ustilago maydis* Rep1 protein is also cleaved by kexin-protease and has a structural role in the fungal cell wall (Wösten *et al.*, 1996; Teertstra *et al.*, 2006). A 11 amino acid long peptide processed by KEX2 in *Cryptococcus neoformans* is required for virulence and to activate the sexual program (Homer *et al.*, 2016; Tian *et al.*, 2018). Since *MiSSP8*-RNAi lines are not impaired in hyphal growth or fungal cell-wall sensitivity, it is unlikely that MiSSP8 or its derived peptides are involved in fungal cell wall structure. In addition, MiSSP8 does not bind to fungal cell wall sugars (data not shown). The precise role of MiSSP8 and its derived peptides in the development of the fungal mantle will require further research.

The fungal proteins identified in our study exhibit the (DW[K/R]R)_n_ motif at their C-termini with a variable number of repetitions but they do not share sequence similarities at their N-termini. Variations in number of tandem repeats is thought to provide functional diversity (Verstrepen *et al*, 2005), suggesting that the identified DW[K/R]R-containing proteins might be involved in various cellular processes and carry out different functions. If KEX2-processed, the variable number of repeats would lead to different number of released peptides. A comparative phylogenomics analysis of 49 fungal genomes revealed that ECM fungi have evolved several times from saprotrophic ancestors, these being either white rot, brown rot or soil decayers, by developing a set of symbiotic genes with rapid turnover, and in particular gain of genes such as MiSSPs (Kohler *et al.*, 2015). Moreover, a recent comparative analysis of five *Amanita* genomes, two ECM symbionts and three asymbiotic species, concluded that several genetic components of the toolkit used by ECM symbionts is already encoded in the genome of their saprotrophic relatives, explaining the recurrent emergence of the ECM symbiosis over time (Hess *et al.*, 2018). Consistent with this finding, MiSSP8 shares similarities with other saprotrophic proteins through its fungal-specific repetitive motif. Since MiSSP8 is likely playing a role in the formation of the pseudoparenchyma of both non-symbiotic (basidiocarp) and symbiotic (ECM) structures, we propose that MiSSP8 function, initially required for *L. bicolor* fruiting body formation, has been recruited for the establishment of the symbiosis. This highlights the dual use of the given secreted protein into symbiotic and non-symbiotic processes. Our analysis also suggest that there are two categories of MiSSPs: the orphan ones, such as MiSSP7 and the ones preexisting in saprotrophic ancestors, such as MiSSP8.

In conclusion, we have characterized the Mycorrhiza induced Small Secreted Protein of 8 kDa (MiSSP8) by combining functional and *in silico* approaches. We demonstrate that MiSSP8 has a functional secretion signal peptide and it contains a repetitive motif containing kexin cleavage sites recognized *in vitro* suggesting the protein might be cleaved in order to become functional. Our data show that MiSSP8 or its derived peptides are decisive factors for symbiosis establishment. The DWRR repetitive motif being also found in SSPs from saprotrophic fungi, we propose that MiSSP8 or its derived peptides could have been initially linked to fruiting body development by participating in pseudoparenchyma formation through fungal hyphae aggregation before being recruited for symbiosis establishment.

## Acknowledgments

This research was sponsored by the Genomic Science Program, US Department of Energy, Office of Science, Biological and Environmental Research as part of the Plant-Microbe Interfaces Scientific Focus Area (http://pmi.ornl.gov) and the Laboratory of Excellence ARBRE (grant no. ANR-11-LABX-0002_ARBRE). The research was supported by the Institut National de la Recherche Agronomique and the University of Lorraine (Ph.D. scholarship to CP and YD). Both Région Lorraine Research council and the European Fund for Regional Development give funding for the Functional Genomics Facilities at Institut National de la Recherche Agronomique-GrandEst. Part of the work was supported by Laboratoire Recherche Sciences Végétales - UPS CNRS, MetaToul (Metabolomics and Fluxomics Facitilies, Toulouse, France, www.metatoul.fr) and the French National infrastructure for metabolomics and fluxomics, www.metabohub.fr, MetaboHUB-ANR-11-INBS-0010. JR was an AgreenSkills Marie Sklodowska Curie postdoctoral fellow co-funded by the European Commission (FP7-267196). We thank Barbara Montanini and Simone Ottonello for sharing the pSUC-modified vectors. We thank Alexandre Kriznik from the Platform of Biophysics and Structural Biology-UMS 2008 IBSLor (CNRS-INSERM-UL) for his technical assistance in circular dichroism and isothermal calorimetry experiments. Funding to A.P. and M.K. was provided by grants from Universidad Nacional de Quilmes, Consejo Nacional de Investigaciones Científicas y Técnicas, and Agencia Nacional de Promoción Científica y Tecnológica, Argentina.

## Authors contributions

CVF and FM designed and managed the project; CP, YD, FG, MK, JR, VP, NFdF, AH, CVF performed the experiments; CP, CVF, MK, AP, VP, NFdF analyzed the data; CP performed the bioinformatic analysis; CP and CVF wrote the manuscript and all authors revised it.

## Legends of Figures

**Figure S1: Fruiting body of *L. bicolor* at two different developmental stages.** The cap and stipe of fruiting bodies from *L. bicolor* were harvested at early and late developmental stages in order to study the expression of *MiSSP8* during the development of the fruiting body.

**Figure S2: *In silico* prediction and experimental validation of MiSSP8 secondary structure**. **(a)** Secondary structure of MiSSP8 as predicted by the JPred 4 server (Drozdetskiy *et al*, 2015). **(b)** Analysis of recombinant MiSSP8 secondary structure by circular dichroism.

**Figure S3: Correlation between protein size sequences and occurrence of the DWRR/DWRR-like motif.** No correlation found between the size of the DW[K/R]R protein sequences identified and the number of motifs they contain (A), even after removing the outliers (B).

**Figure S4: Spectrum obtained for the peptide DSDWR in LC-MS/MS.** The selected precursor ion (*m/z* 678.6) is corresponding to the peptide DSDWR (M+H)^+^; after fragmentation (CE 40V), y ions are observed as predicted in Protein Prospector (*m/z* 175.1; 361.1; 476.2; 563.2; 678.2).

**Figure S5. *L. bicolor missp8* RNAi mutants are not impaired in their saprotrophic growth in both control medium (P5) or under cell-wall stress conditions.** Viability of *MiSSP8* RNAi lines was assessed by growing the different strains used (Wild-type (WT), empty vector control (ev7) and the four RNAi lines generated on either control medium (P5) or under cell-wall stress condition (Congo Red).

**Supplementary Table S1**: List of genomes scanned and proteins sequences containing from two to twenty repetitions of the DW[K/R]R motif.

**Supplementary Table S2**: Molecular analysis of *Laccaria bicolor MiSSP8*-RNAi lines. Molecular analysis of the four RNAi lines generated expressing RNAi hairpin targeting the MiSSP8 encoding transcript.

## Notes

#### Summary of Updates

The repetitive motif of MiSSP8 located at the C-terminal part of the protein contains a kexin cleavage site recognized in vitro. Therefore, the Figure 5 showing in planta localization of the full length protein has been replaced by a new Figure 5 showing the cleavage of MiSSP8 into several small peptides. The text has been modified accordingly.

