## Supplementary figures and images for "*Laccaria bicolor* MiSSP8 is a small-secreted protein decisive for the establishment of the ectomycorrhizal symbiosis"

### Supplemental Figure S1

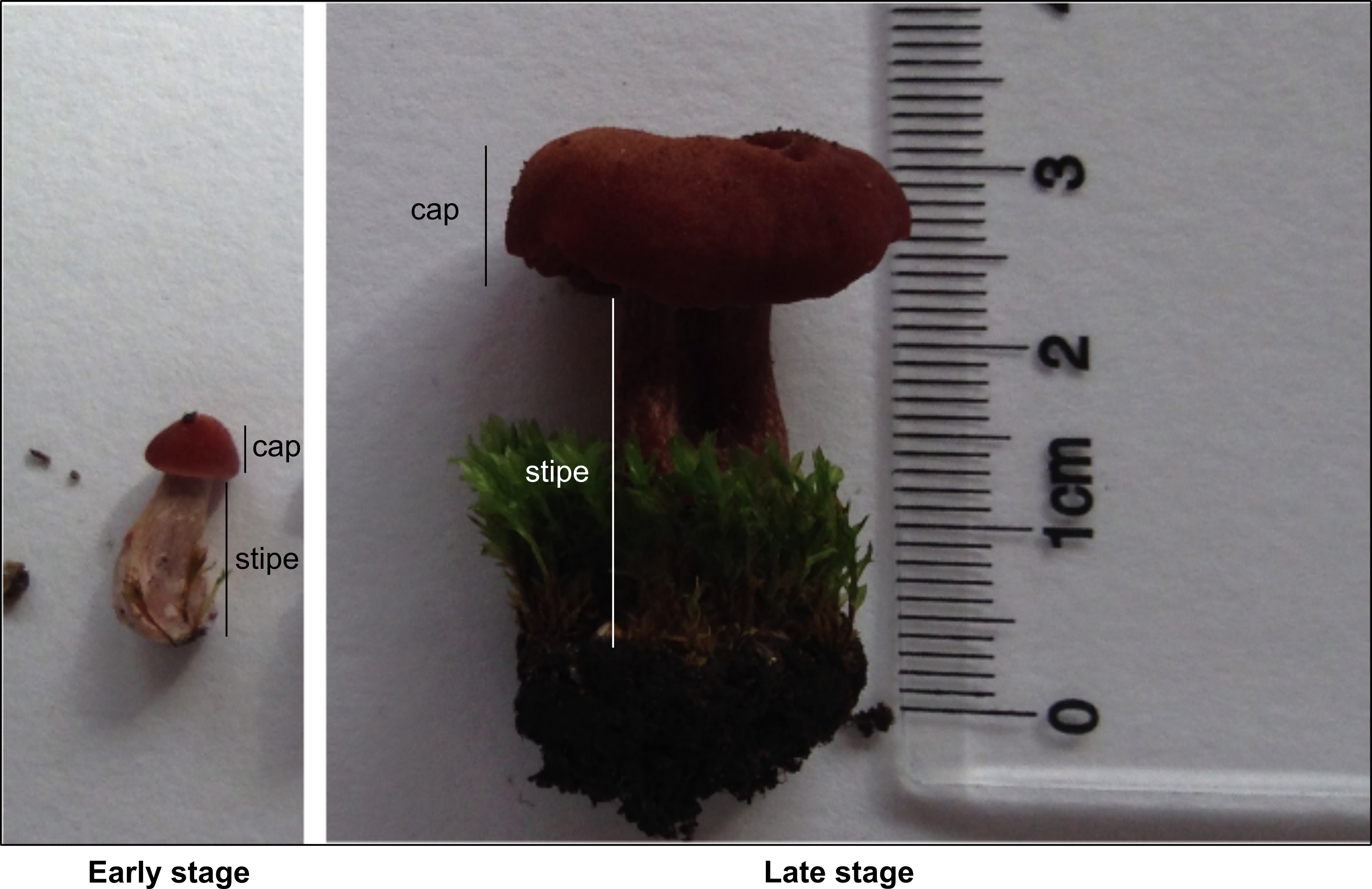

### Supplemental Figure S2

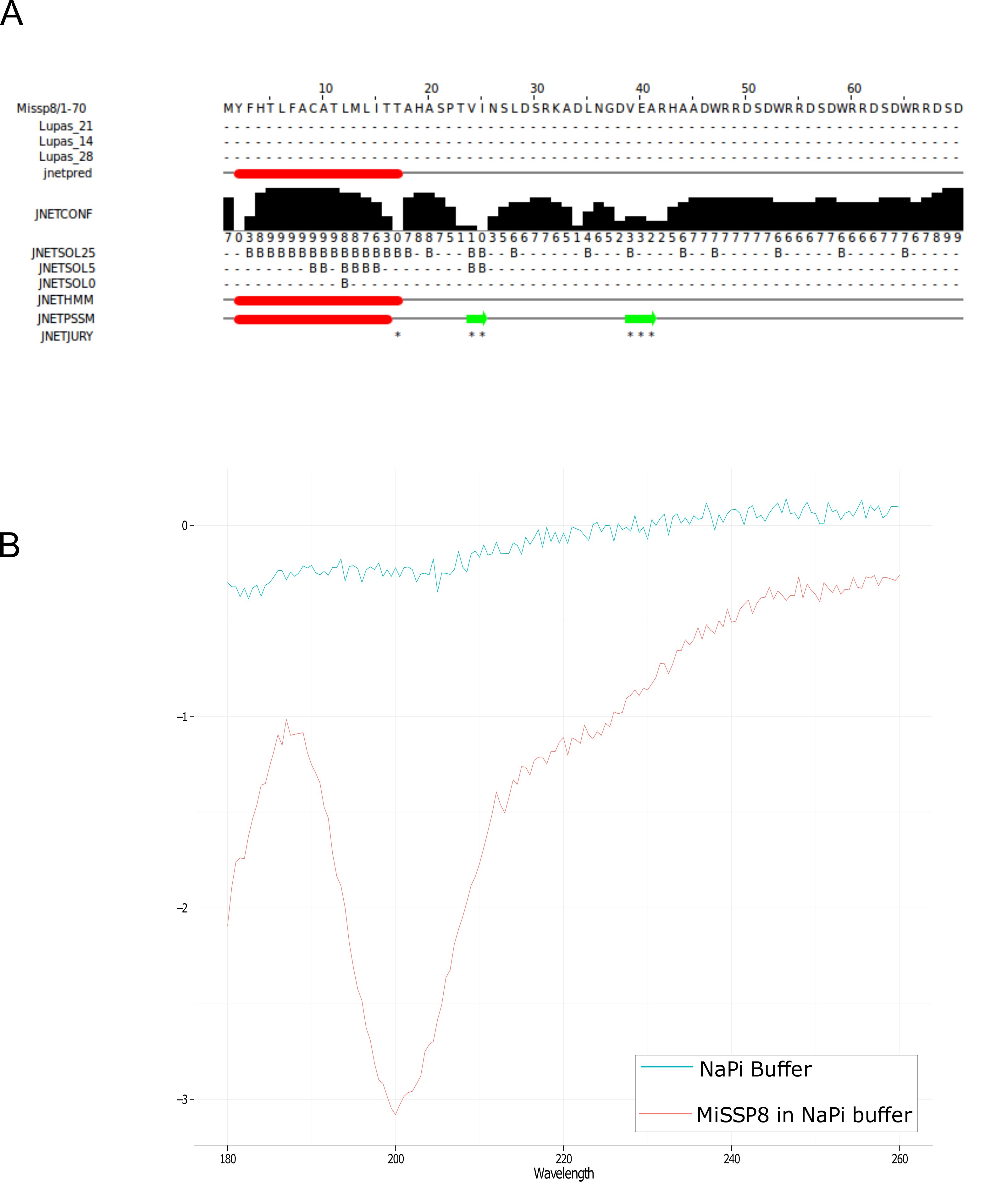

### Supplemental Figure S3

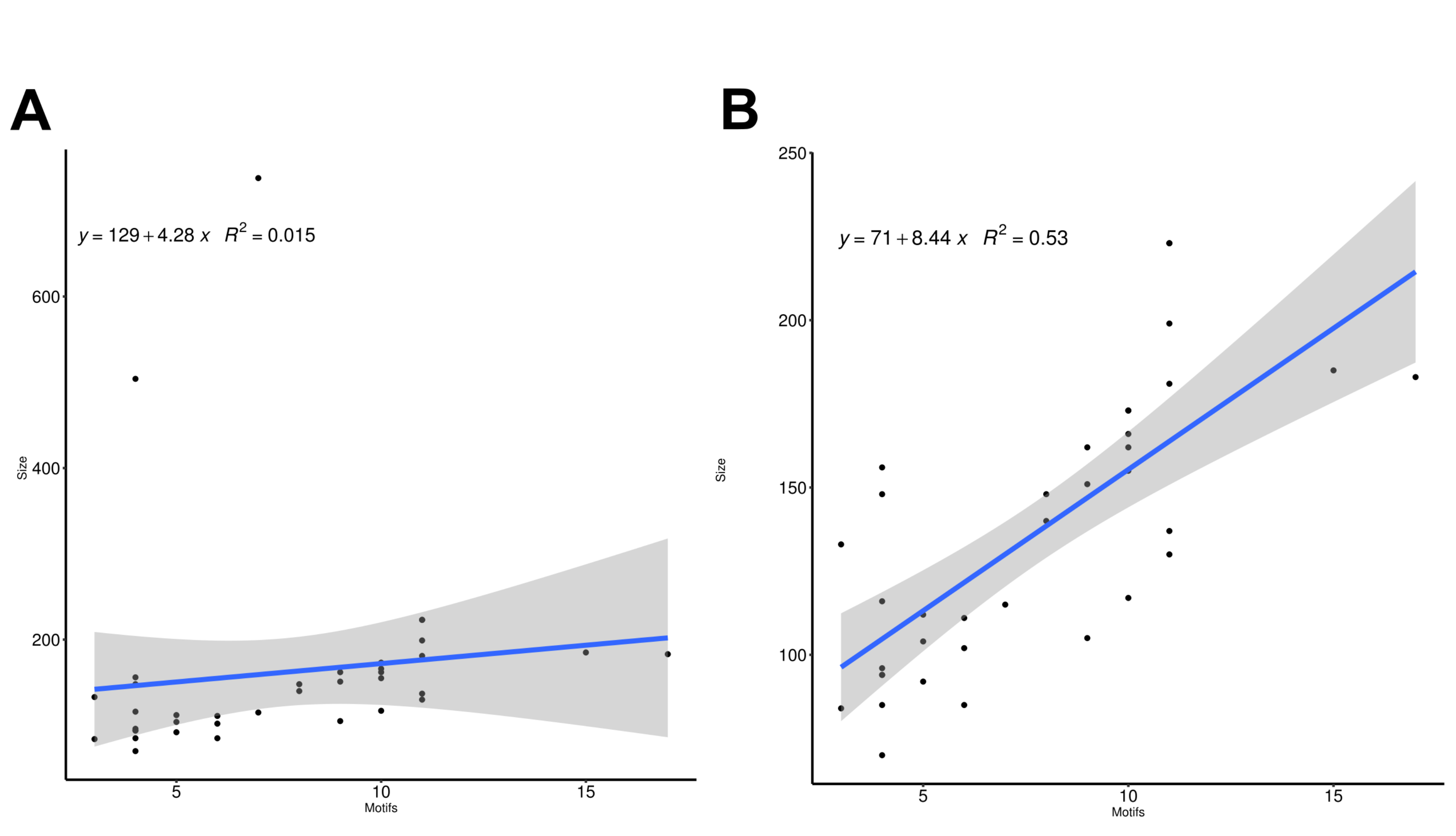

### Supplemental Figure S4

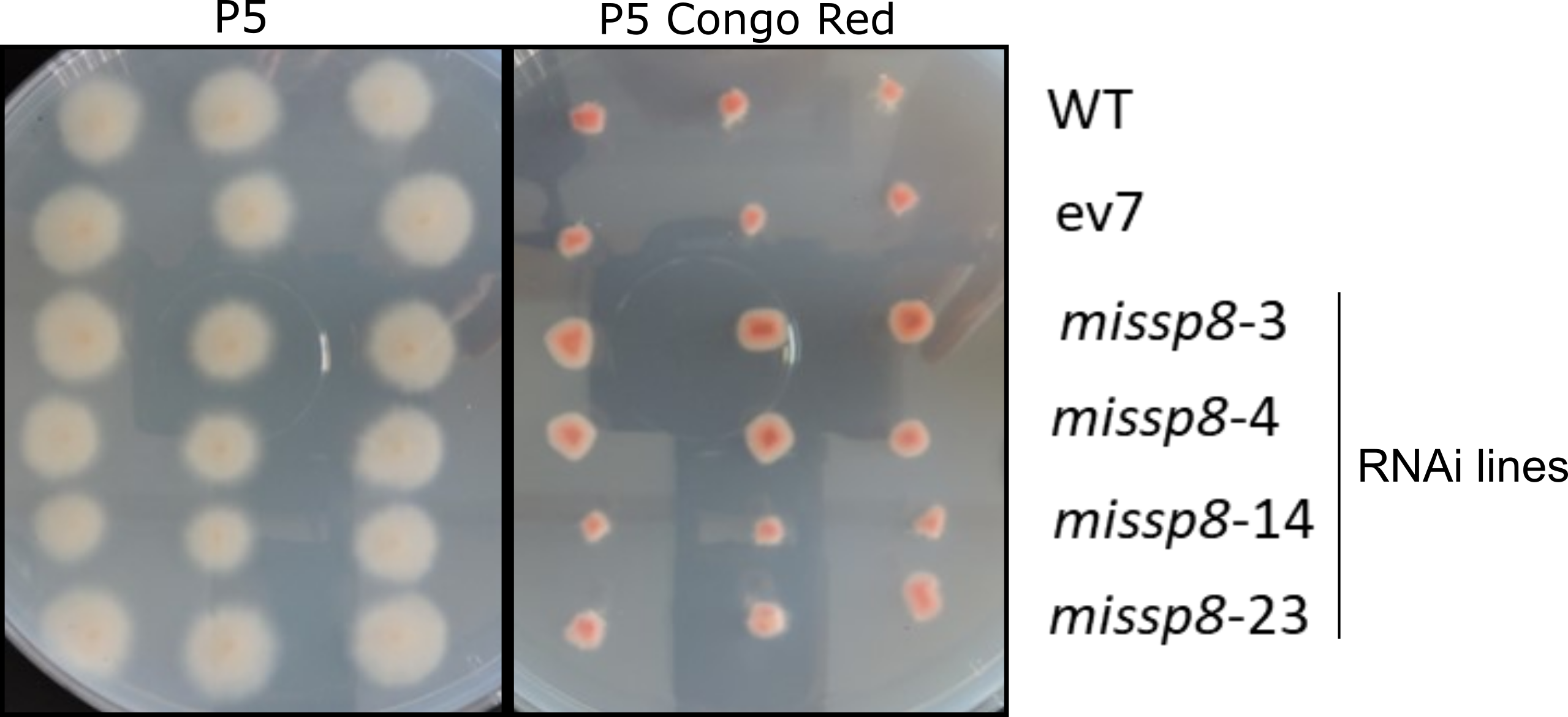

### Supplemental Figure S5

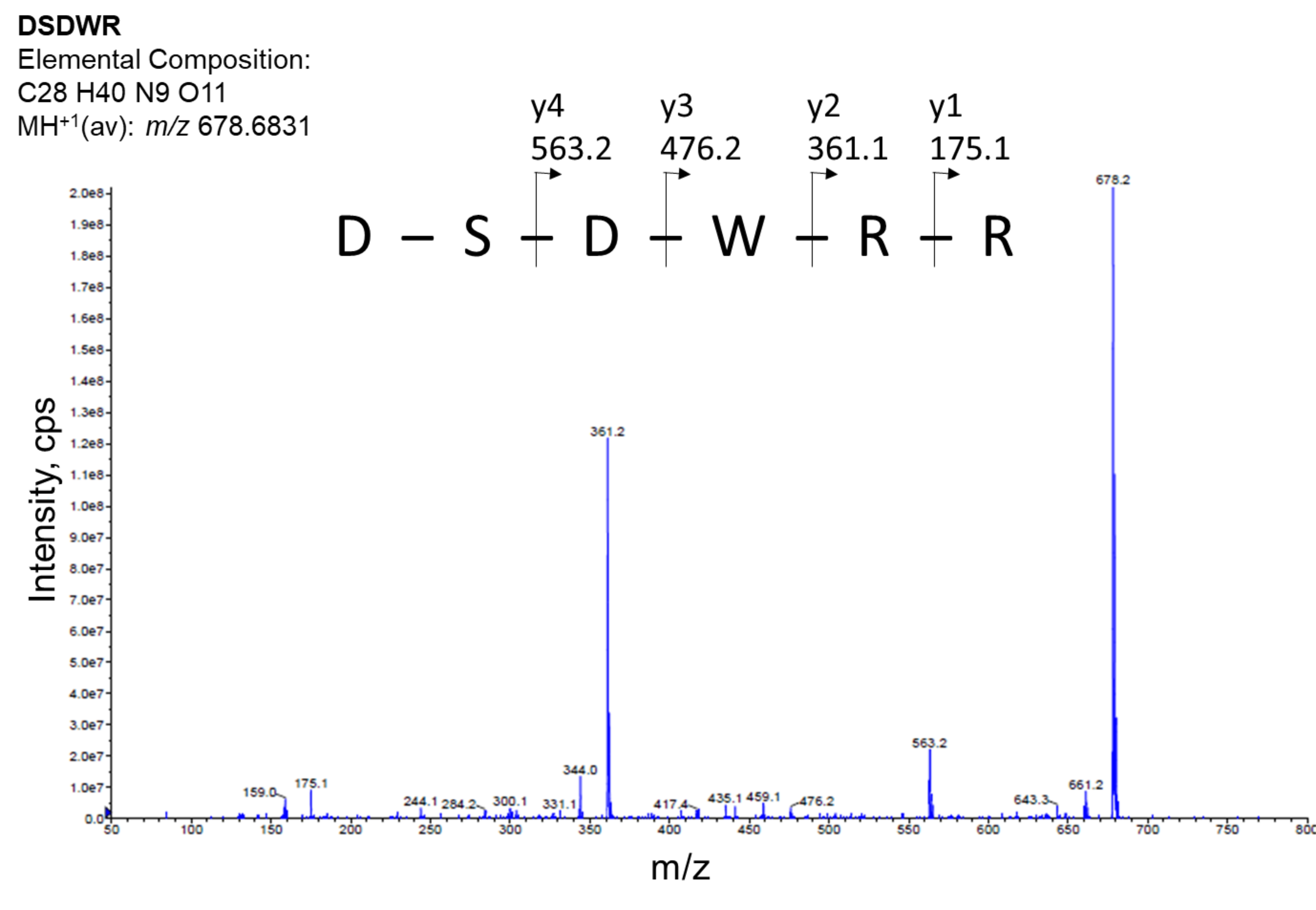
